# Using seedling phenotypic traits to select local seed sources for large-scale restoration: methods and outcomes in a Great Basin case study

**DOI:** 10.1101/2020.04.10.036459

**Authors:** Elizabeth A. Leger, Sarah Barga, Alison C. Agneray, Owen Baughman, Robert Burton, Mark Williams

## Abstract

Establishing plants from seed is often a limitation to restoration success in semi-arid systems. For restoration purposes, managers can either use widely-available commercial seeds, which are often sourced from far outside the seeding area, or take extra steps to use locally collected seeds. If local seeds have traits more conducive to seedling establishment in degraded sites, they could increase restoration success. Here, we asked whether wild-collected seeds of two native perennial grasses, *Elymus elymoides* and *Poa secunda*, had more favorable characteristics than commercial sources. Seeds were collected from four populations within the Winnemucca District of the Nevada Bureau of Land Management, which manages lands within the Great Basin, US. Collections were screened for seed and seedling characteristics associated with increased plant performance in invaded Great Basin systems, and we provide a detailed methodology for these measurements. Relative to commercial seeds, wild-collected seeds had more characteristics identified as beneficial for seedling establishment including earlier emergence, higher specific root length, more root tips, and smaller overall size (*E. elymoides*), and earlier emergence, longer roots, higher root mass ratio, and more root tips (*P. secunda*). Commercial sources had significantly larger seeds than wild populations, a trait that had mixed effects on performance in previous research, and one that may change as wild collections are increased in agronomic conditions. These results suggest that locally-sourced populations are more likely to perform well in invaded areas, providing support for efforts to collect, screen, and increase local sources of seeds to improve restoration success.

**Implications for Practice:**

- Collecting and increasing local seeds requires more time and effort than purchasing commercially-available seeds, but if these sources have a greater chance at surviving in restoration sites, this effort is warranted
- In our study, remnant local populations possessed more potentially adaptive traits than commercially-available alternatives, indicating they may be superior sources for the restoration of disturbed sites in their local regions
- Screening multiple seed sources for potentially adaptive seed and seedling traits can be a relatively quick and effective way to select the most promising seeds for increase

## Introduction

Selecting seeds for restoration is an important step, as incorrect seed selection can set a project up for failure before it even begins. While evidence for restoration-relevant concepts like local adaptation is abundant (Leimu & Fischer 2008; Hereford 2009; Baughman et al. 2019), applying this knowledge to restoration practices at management scales can present challenges (Vander Mijnsbrugge et al. 2010; Gibson et al. 2016), and seed used for restoration is often selected for reasons other than optimal performance in a particular habitat (Davies et al. 2011; Ladouceur et al. 2018). For example, in the Great Basin, common garden studies have identified large-scale patterns of local adaptation across the range of widespread species (Baughman et al. 2019), and seed transfer zones have been described for many common species (e.g., Bower et al. 2014). Despite this readily available information, most seed used for restoration and rehabilitation in this region are either cultivars or populations selected for characteristics that may differ from priorities of restoration practitioners (Leger & Baughman 2015). In many cases, seeds are sourced from geographic locations well outside the regions where seeding is occurring (Jones & Larson 2005; Pilliod et al. 2017).

A primary reason for the reliance on non-local seeds is efficiency in seed production and economies of scale (Chivers et al. 2016), but managers can collect and increase seeds from local areas, even though this may take extra effort and time. This effort is more common for smaller-scale restoration (e.g. Houseal & Smith 2000; Kiehl et al. 2010), but it is even possible for restoring large areas of public lands in the Western US, where some of the largest seeding projects are regular occurrences (Pilliod et al. 2017). For example, for federal management agencies it is possible to commission the large-scale production of locally-collected seeds directly from growers via a type of contract called indefinite delivery/indefinite quantity or IDIQ (Riley et al. 2015). This type of contract allows for lower-risk production of seeds from untested, uncommon, or potentially challenging species or populations by seed producers, providing flexibility in seed availability in particular regions (Peppin et al. 2010). While this strategy requires greater up-front investment, its flexibility allows managers to take advantage of remnant populations of native species persisting in disturbed and invaded areas, which could be superior at establishing in typical restoration scenarios (Leger 2008; Goergen et al. 2011).

Here, we present an example of a collaboration between researchers and federal land managers designed to assess the potential for increased restoration success with local seeds. Our goal was to collect seeds from multiple wild populations of native perennial grasses within a particular management area, gather sources of widely-available cultivars and commercial varieties, and screen these seed sources to identify which had the greatest number of phenotypic traits associated with increased seedling performance in disturbed, invaded restoration scenarios (hereafter, “potentially adaptive traits”; Appendix 1). We focused on two species, *Elymus elymoides* (Raf.) Swezey and *Poa secunda* J. Presl, because they are widespread throughout the management area, are particularly good at persisting in disturbed and invaded environments, and are commonly seeded in projects in this region (Booth & Grime 2003; Goergen et al. 2011; Stevens et al. 2014; Leger & Baughman 2015).

Here, we also describe the process of identifying potentially adaptive traits (Box 1), and methods for measuring seed and seedling traits (Box 2) for grasses and other species (Box 3). Our project benefited from previous experiments that focused on identifying the potentially adaptive traits associated with seedling performance of *E. elymoides* and *P. secunda* in the presence of the highly competitive exotic annual grass *Bromus tectorum* L. Through a series of greenhouse and field studies (Appendix 1), we have found a number of traits associated with greater performance or survival of plants in these systems including: earlier emergence, longer roots, higher root mass ratio (RMR; the amount of total biomass allocated to roots), a larger number of root tips, and higher specific root length (SRL; a measure of root length to root weight, with higher values typically indicating greater allocation to fine roots), and smaller seedling size (for *E. elymoides*) (Leger & Baughman 2015; Leger & Goergen 2017; Leger et al. 2019). Seed size has also been important, though the relationship between seed size and success has had mixed outcomes, with larger seeds performing better in some situations, and smaller seeded plants performing better in others (e.g., Kulpa & Leger 2013; Leger et al. 2019). We expected that locally-collected populations from disturbed sites would possess a greater number of potentially adaptive traits than widely-available commercial varieties, indicating that there would be value in expending the effort to increase them for use in local restoration.

### Box 1. Determining important traits in your study system

In restoration projects, the ultimate measure of a planted seed’s success is its ability to grow and survive in its new habitat. Predicting establishment success can be improved by research efforts that identify potentially adaptive phenotypic trait values in seeds and seedlings. There are multiple ways to identify which phenotypic traits are important in your study system, including the following:

1. **Review the literature** for published papers that have conducted common garden studies with your species of interest, and observe which phenotypic traits have the strongest correlations with environmental variables. For example, in a common garden study of *Achnatherum hymenoides* (Roem. & Schult.) Barkworth, traits associated with flowering phenology, leaf abundance, and leaf morphology, were correlated with annual temperature and precipitation at the collection location (Johnson et al. 2012). These would be an excellent starting place for screening, looking for populations that have trait values closest to the conditions at your restoration site.
2. **Infer likely important traits** from a meta-analysis of cross-species patterns. If no work has been done on your species of interest, one could take a best-guess based on traits important for similar functional groups. For example, in a review of what is known about local adaptation in the Great Basin, traits like leaf length and flowering phenology were frequently correlated with environmental variables (Baughman et al. 2019).
3. **Expert opinion, ecological paradigms**. The adaptive traits we have identified in invaded Great Basin systems (e.g., early emergence, increased root allocation, etc.) may not be adaptive in other ecosystems experiencing, for example, greater precipitation, different types of invasive species, or different forms of herbivory. In cases where no information is available, general information about likely adaptive characteristics (investment in shoot growth, defensive compounds, etc.) could be inferred by studies on, for example, competitive dynamics or plant/herbivore interactions in your system.
4. **Direct tests**. Conducting experiments comparing the distribution of phenotypic traits before/after restoration can directly determine the strength and direction of natural selection on particular traits (Kulpa & Leger 2013), or greenhouse/field experiments where you can link success with independently-measured plant phenotypes (e.g. Rowe & Leger 2011; Leger et al. 2019).

**Box 1, Figure 1.**
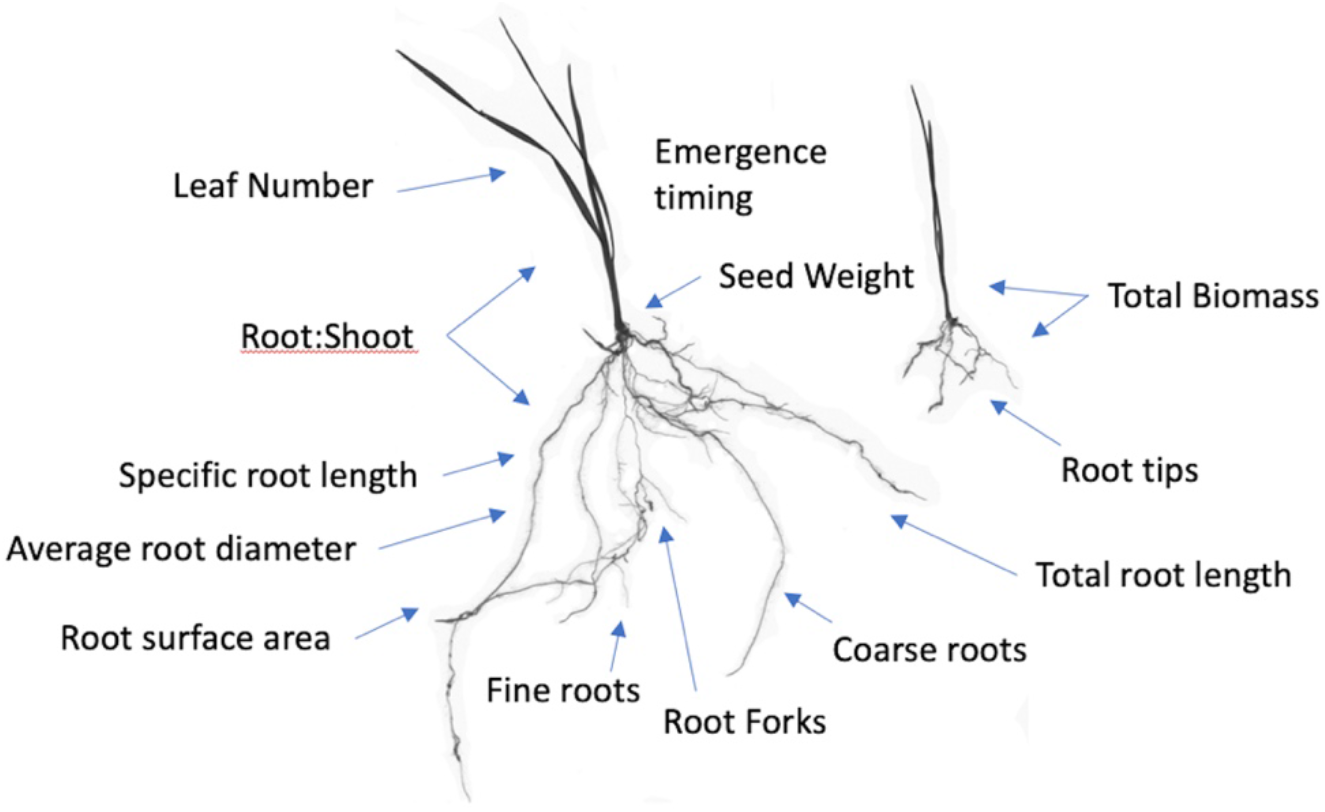
Images of two *Poa secunda* plants harvested 35 days after emergence in the greenhouse, along with examples of the phenotypic traits that can be measured on plants of this size. Through experiments, we have determined that root characteristics of very young plants can be correlated with seedling performance in competitive, moisture limited conditions.

### Box 2. Working with seedling roots: tips and methods

Screening seedlings for phenotypic traits requires technique and practice. To avoid some common pitfalls, we are sharing our tips and successful methods:

- Choose a growing medium that is appropriate for your system (e.g., native topsoil with little organic matter works well for Great Basin plants) and mix 50:50 with an inert material (washed decomposed granite or sand) to ease soil removal. Soils with organic matter or vermiculite become highly problematic during root screening, as it is difficult to separate from roots.
- We used perlite as the inert material in this experiment, but we don’t recommend it or use it any longer, as roots can grow into the perlite, and it has a tendency to rise to the top of pots, obscuring the soil surface.
- Plant seeds directly in pots larger than your anticipated root structures will grow in the allotted time
- To track emergence timing, monitor the soil surface daily for new cotyledons and track individual plant pot emergence
- Maintain a consistent age for harvesting the seedlings (e.g., 10 or 15 days after emergence for quickly growing grasses or forbs and 35 or 50 days for slower-growing species). Ideally, the developmental stage where phenotypic traits have the strongest impact on seed survival would be determined from experiments, but if not, consider selecting more than one stage for screening
- Consider planting in staggered blocks, with at least one of each seed source planted in each block. With one person, we found that processing around forty pots per day was realistic.
- When harvesting, soak the entire pot in a tub of water to loosen the soil, gently tap the pot against the sidewalls of the tub until roots are completely released. Next, transfer the entire plant into a series of cleaner tubs until the seedling is free of debris.
- Do not rub or pick at the roots, since fine roots can be easily lost in this stage.
- Use the WinRhizo software to directly scan your root images, avoiding the perils of file conversion and nonconforming pixel aspect ratios associated with scanning with other software.
- During the screening process, roots are suspended in water, which makes them easy to arrange. If bubbles or particulates are obscuring roots in images, we find that using water that has sat out for 24 hours or filtered water results in cleaner images.

**Box 2, Figure 2.**
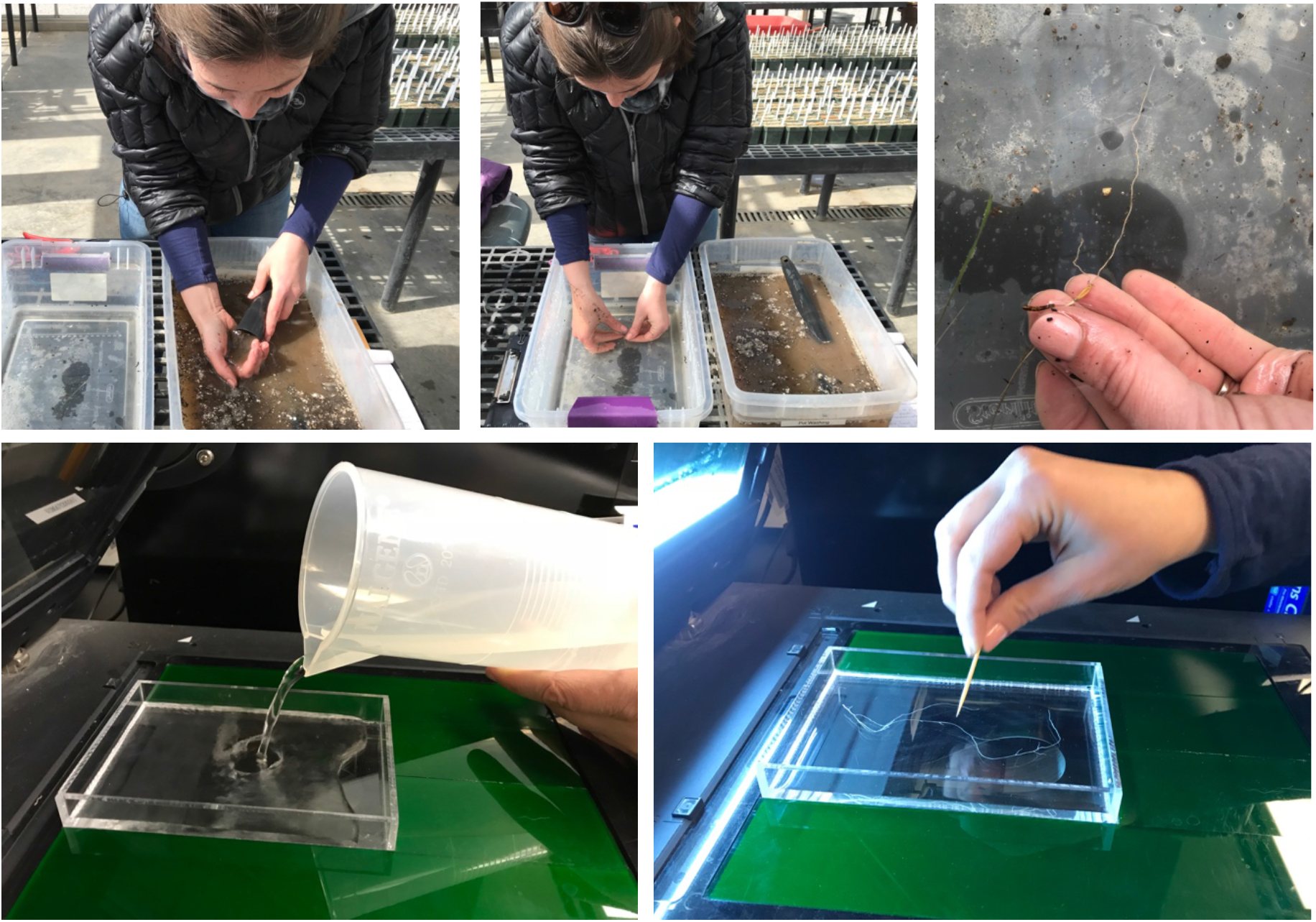
Clockwise from top-left: After a brief soak, removing seedling and soil from planting tube. Second rinse in clear water to carefully remove soil particles. A clean root, ready to separate from shoot for scanning/drying/weighing. Preparing to scan roots using the WinRhizo system. Arranging roots to reduce overlap for an accurate scan.

### Box 3. Considerations for root screening for different functional groups

Each species and functional group are unique and may provide different challenges. The most notable factors impacting seedling trait screening that we have observed are variance in 1) emergence timing and seed dormancy, 2) root structure and elasticity, and 3) growth rate.

**Emergence timing, seed dormancy**. There may be significant variation in emergence timing between species and even populations. For instance, we have found that while all populations of *Elymus elymoides* reliably emerge within 20 days of planting, *Achnatherum thurberianum*, another native perennial grass, took anywhere between 6 and 166 days to emerge, with large differences among populations. Other species can exhibit dormancy, with differences among populations (e.g., Barga et al. 2017), which would need to be addressed before screening. Thus, even screening for characteristics of young (10-days post-emergence, for example) seedlings can take many months to complete, if emergence timing is delayed.
**Root structure and elasticity**. Particularly between functional groups, the size, resilience, and general architecture of the roots will vary, having cascading impacts on the difficulty of cleaning roots for scanning and the resulting accuracy of the measurements. For instance, roots of perennial grasses tend to be easily cleaned and measured, since they are elastic, resistant to breaking, and lay flat with relative ease. Alternatively, older shrub roots tend to be brittle and resist lying flat for root measurements, so they require greater care and effort during the cleaning and scanning process. It is still possible to scan broken roots and measure some characters (length, diameter), but root tips should not be assessed on broken roots.
**Growth rate**. The speed at which each species grows will vary significantly, and we recommend a pre-screening trial to assess how fast your species of interest will grow to help determine the appropriate harvest date. When choosing your harvest date, if you don’t have prior information on what stage is most critical in restoration, we suggest selecting an age where seedlings are starting to show obvious differentiation among populations (e.g., different number of leaves or varying root length), but are not yet outgrowing their containers.

**Box 3, Figure 3.**
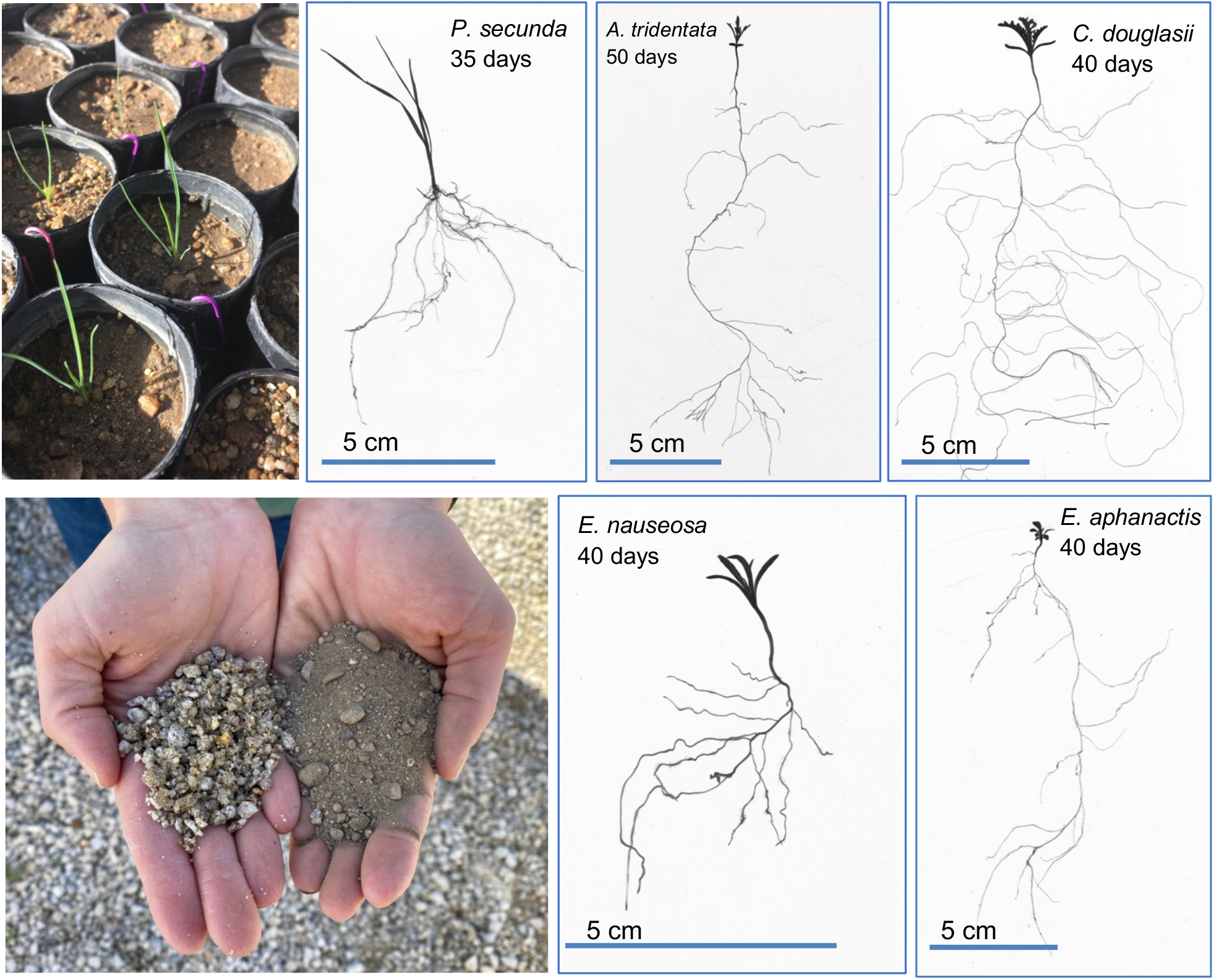
Top left: Above-ground tissue of *Poa secunda* before harvest. Bottom left: Examples of field soil and decomposed granite, before mixing. All other panels: Plant and root scans of grasses (*Poa secunda*), shrubs (*Artemisia tridentata, Ericameria nauseosa*), and forbs (*Chaenactis douglasii, Erigeron aphanactis*) of various ages.

## Methods

Seeds from two wild populations each of *E. elymoides* and *P. secunda* were collected from remnant native populations by a Seeds of Success (SOS) seed collection team, a national program run by the Bureau of Land Management (BLM) designed to collect and store native seeds for restoration (Haidet & Olwell 2015). These populations originated from within low elevation, disturbed areas of the Great Basin (Table 1). Sites were located within the boundaries of the BLM’s Winnemucca District, which includes 3.3 million hectares of lands, many of which experience frequent fires, significant invasion of exotic species, and extensive reseeding efforts (Pilliod et al. 2017). At all four collection sites, target species were co-occurring with an abundance of invasive annual weeds (*B. tectorum, Ceratocephala testiculata* Crantz) Roth, *Chorispora tenella* (Pall.) DC., *Descurainia sophia* (L.) Webb ex Prantl, *Erodium cicutarium* (L.) L’Hér. ex Aiton), *Lepidium perfoliatum* L., *Salsola tragus* L., *Sisymbrium altissimum* L.), and sites showed evidence of grazing use and a history of wildfire. Mean annual precipitation at wild collection sites varied, but estimates of 30-year normal average precipitation were less than 270 mm/year at all sites (PRISM Climate Group, http://www.prism.oregonstate.edu; Grass Valley *E. elymoides:* 268 mm, Sonoma *E. elymoides*: 243 mm, Button Point *P. secunda*: 268 mm, Winnemucca Mt. *P. secunda*: 198 mm). Seeds were collected by SOS teams in June and July of 2014 and 2015, with a bulk collection from ~1000 (*E. elymoides*) or ~10,000 (*P. secunda*) plants. In addition, four commercially-available seed sources of seed were screened, including Hanford and Mt. Home *P. secunda*, and Vale and Toe Jam Creek *E. elymoides*, which were provided by L&H Seeds or the Winnemucca BLM district. These sources represent the most proximate or climatically appropriate of the widely-available commercial sources for use in the portions of this district in need of restoration.

**Table 1.** Species and sources of seeds screened, as well as mean values and standard errors (in parentheses) for seven potentially adaptive seed and seedling traits. Superscript letters are presented when ANOVA indicated significant differences among source populations, and values with different letters are significantly different from each other based on Tukey’s HSD tests, with a P <0.05 threshold for determining significance. Bold and shaded cells indicate seed sources with significantly favorable values for each trait. Comparisons were made among ELEL collections and POSE collections separately.

| Species | Source | Type | SOS or citation<sup>1</sup> | Seed wt. (mg) | Days to emergence | Root length (cm) | RMR | # Root tips | SRL | Total size (mg) |
| --- | --- | --- | --- | --- | --- | --- | --- | --- | --- | --- |
| ELEL | Grass Valley | Wild | NV020-7 | 2.00<sup>c</sup><br>(0.02) | 10.3<sup>ab</sup><br>(0.6) | 12.7<br>(0.7) | 0.52<br>(0.01) | 84.9<sup>ab</sup><br>(8.1) | **98.4<sup>a</sup>**<br>**(4.7)** | **2.5<sup>c</sup>**<br>**(0.08)** |
| ELEL | Sonoma | Wild | NV020-1 | 2.20<sup>b</sup><br>(0.02) | **9.3<sup>b</sup>**<br>**(0.6)** | 14.4<br>(0.9) | 0.50<br>(0.02) | **99.9<sup>a</sup>**<br>**(9.1)** | **98.4<sup>a</sup>**<br>**(4.6)** | **2.9<sup>c</sup>**<br>**(0.07)** |
| ELEL | Toe Jam Creek | Commercial | Jones et al. 2011 | **2.56<sup>a</sup>**<br>**(0.07)** | 11.0<sup>ab</sup><br>(0.5) | 12.9<br>(1.0) | 0.52<br>(0.01) | 69.0<sup>b</sup><br>(7.5) | 70.9<sup>b</sup><br>(4.7) | 3.6<sup>a</sup><br>(0.16) |
| ELEL | Vale | Commercial | Jones 2014 | **2.60<sup>a</sup>**<br>**(0.05)** | 11.1<sup>a</sup><br>(0.7) | 12.8<br>(1.5) | 0.56<br>(0.02) | 81.6<sup>b</sup><br>(14.9) | 78.1<sup>b</sup><br>(7.4) | 2.8<sup>bc</sup><br>(0.18) |
| POSE | Button Pt. | Wild | NV020-2 | 0.44<sup>ab</sup><br>(0.02) | **11.7<sup>b</sup>**<br>**(0.4)** | **7.3<sup>a</sup>**<br>**(0.4)** | **0.62<sup>a</sup>**<br>**(0.01)** | **38.9<sup>a</sup>**<br>**(2.9)** | 126.6<br>(5.4) | 0.95<sup>a</sup><br>(0.04) |
| POSE | Winn. Mt. | Wild | NV020-4 | **0.45<sup>a</sup>**<br>**(0.01)** | 13.7<sup>a</sup><br>(0.6) | 5.2<sup>b</sup><br>(0.3) | **0.65<sup>a</sup>**<br>**(0.02)** | 27.3<sup>b</sup><br>(2.4) | 128.1<br>(8.2) | 0.70<sup>b</sup><br>(0.04) |
| POSE | Hanford | Commercial | Majerus et al. 2009 | **0.45<sup>a</sup>**<br>**(0.02)** | 12.3<sup>ab</sup><br>(0.5) | 6.0<sup>ab</sup><br>(0.4) | 0.56<sup>b</sup><br>(0.01) | 32.2<sup>ab</sup><br>(2.5) | 138.5<br>(6.2) | 0.79<sup>ab</sup><br>(0.03) |
| POSE | Mt.<br>Home | Comme<br>rcial | Lambert<br>et al<br>2011 | 0.40<sup>b</sup><br>(0.01) | 11.8<sup>ab</sup><br>(0.6) | 6.6<sup>ab</sup><br>(0.6) | 0.60<sup>ab</sup><br>(0.02) | 36.3<sup>ab</sup><br>(4.3) | 124.3<br>(11.4) | 0.94<sup>a</sup><br>(0.09) |

<sup>1</sup>The SOS seed collection number or the citation describing the release of named seed sources is shown, if available; Hanford *P. secunda* and Vale *E. elymoides* are unregistered, wild-collected, agriculturally-increased seed originally collected near Hanford, WA and Vale, OR, respectively.

For each seed source, seeds were weighed in 10 batches of 20 seeds, and values were used to calculate average seed weights. We planted 90 seeds per seed source in containers (Stuewe and Sons SC10 3.8 cm x 21 cm cone-tainers), in a 50:50 topsoil and perlite mixture. We planted in a randomized block design over nine planting periods to allow for a staggered harvest. Planting began on October 21, 2015 and continued every Monday, Wednesday, and Friday until November 9^th^, with 10 seeds per seed source planted every day. Seeds were watered daily, and we noted the day each seedling emerged. Plants were harvested ten days after emergence, at which point we carefully washed the roots from the soil (Box 2). Roots and shoots were separated, and root samples were scanned and analyzed using WinRhizo software (Regents Instruments, Siante-Foy, QC, Canada). Roots and shoots were then dried and weighed separately for each plant. From WinRhizo, we gathered the length of each root system and the number of root tips. RMR was calculated as root biomass/total biomass, and SRL was calculated as root length in m/root weight in g.

Data were analyzed using analysis of variance (ANOVA) with the program R (R Core Team 2019). Each response variable was log transformed and scaled as needed to meet model assumptions, with seed source as a fixed factor in each model. Models with significant seed source effects were followed by Tukey’s HSD tests to determine significant differences among sources. For each phenotypic trait, we identified the population(s) with characteristics most likely to be adaptive in scenarios where native species are seeded into disturbed, invaded systems (Appendix 1), and tabulated the number of potentially adaptive traits for each seed source.

## Results

### Seed size

Seed weights of *E. elymoides* differed among populations (F_3,36_ = 44.0, *P* < 0.0001). Commercially-available, field increased *E. elymoides* sources (Vale, Toe Jam Creek) were significantly heavier than either population of wild-collected seeds, and the Grass Valley population had the smallest seeds (Table 1). Seed sizes were more similar among *P. secunda* sources, but there were still significant differences among populations ((F_3,36_ =3.0, *P* =0.0412). Seeds from the Mt. Home source were significantly smaller than those from the Hanford and Winnemucca Mt. sources; Button Pt. had intermediate values.

### Days to emergence

There were significant differences among *E. elymoides* populations (F_3,330_=3.0, *P* = 0.0313). The Sonoma wild collection of *E. elymoides* emerged significantly earlier than other *E. elymoides* seed sources, followed by the Grass Valley wild collection, Toe Jam Creek, and Vale. There were also significant differences among *P. secunda* sources in emergence timing (F_3,213_=3.5, *P* = 0.0171), with the Button Pt. population emerging earliest, followed by Mt. Home.

### Root length

There were no significant differences in 10-day root length among *E. elymoides* collections (F_3,235_=1.1 *P* = 0.3386). There were significant differences among the *P. secunda* collections (F_3,213_ =3.8, *P* = 0.0106): Button Pt. had the longest roots, followed by Mt. Home, Hanford, and Winnemucca Mt.

### Root mass ratio (RMR)

Root allocation was very similar among *E. elymoides* sources (no significant differences among populations, F_3,235_=1.9 *P* = 0.125), and was around 50% of total biomass for most sources. Allocation to roots was generally higher in *P. secunda*, with values over 60% for all but the Hanford seed source. *Poa secunda* populations differed in root allocation (F_3,211_=6.6, *P*= 0.0003); the two wild *P. secunda* collections had the highest root allocation, with Winnemucca Mt. the highest, followed by Button Pt. and Mt. Home.

### Root tips

*Elymus elymoides* populations differed in root tip production (F_3,235_=2.8 *P* = 0.0417), with the Sonoma population having the most root tips, followed by the Grass Valley population. There were significant differences among *P. secunda* seed sources (F_3,211_=2.9, *P* = 0.0347), with the Button Pt. collection having the most root tips, followed by Mt. Home, Hanford, and Winnemucca Mt.

### Specific root length (SRL)

SRL was significantly higher for the two wild-collected *E. elymoides* populations (F_3,235_=8.5, *P* < 0.0001). While SRL was overall higher for *P. secunda* collections relative to *E. elymoides*, *P. secunda* collections did not differ significantly in this measure (F_3,211_=0.7, *P* = 0.5730).

### Total size

There were strong differences in size among *E. elymoides* populations (F_3,235_=10.4, *P* <0.0001). The Toe Jam Creek *E. elymoides* source produced the largest plants, and the Grass Valley and Sonoma wild collections were the smallest. *Poa secunda* plants were much smaller overall, and there were significant differences among *P. secunda* collections in overall size (F_3,211_=5.8, *P* = 0.0008). The Button Pt. and Mt. Home collections were the largest, followed by the Hanford and Winnemucca Mt. collections. Note that we did not highlight any particular size as a potentially adaptive characteristic for *P. secunda*, as previous work did not identify size as an important predictor of survival in this small-statured species.

Overall, when highlighting the traits, we assumed to be adaptive in *E. elymoides* seed sources, the Sonoma wild population has the greatest number of potentially adaptive traits (4 out of 7 measured traits), followed by the Grass Valley wild population (2 traits). For *P. secunda* seed sources, the wild collections again had the greatest number of potentially adaptive traits, with Button Point (4 out of 6 considered traits) and Winnemucca Mt. (2) well ahead of the two commercial collections (0, 1).

## Discussion

There are many barriers to developing restoration strategies that merge the best scientific information with on-the-ground practices (Enquist et al. 2017). Here, we present a collaborative project designed to identify potentially adaptive phenotypic traits within multiple possible seed sources for use in restoration in a western landscape. Focusing on two perennial grass species commonly seeded in the Great Basin, we found variation among collections for a suite of traits that we previously found to be adaptive in field and greenhouse studies. Based on our screening of seed and seedling traits, we predict that the wild-collected Sonoma *E. elymoides* and the Button Pt. *P. secunda* would be the most promising seed sources of those evaluated for establishing in semi-arid, invaded Great Basin sites. The commercially-available seed sources scored higher than field-collected seeds for one potentially adaptive trait, seed size, which was larger overall in commercially-produced seeds. In all other cases, when there were significant differences among collections in potentially adaptive traits, the wild sources had trait values in the direction that our previous research suggests would be adaptive in field plantings (Appendix 1).

Seed size is a trait known to be affected by maternal growing environment (Roach & Wulff 1987), so it is difficult to directly compare this trait among agriculturally-produced plants and field-collected plants. Wild-collected seeds may well increase in size when grown under more resource-rich conditions, or there may be genetic differences in seed size between the wild collections and cultivars; common garden studies would be needed to differentiate between these possibilities. We had an opportunity to weigh agriculturally-increased seeds of Button Pt. *P. secunda* and Sonoma *E. elymoides* after our recommendations lead to selection of these sources for increase under a BLM IDIQ contract, and seed weights for both of these populations increased only slightly: the average seed size of Button Pt seeds from the wild was 0.44 mg, and field increased seeds had an average seed weight of 0.46 mg. The same was true for Sonoma *E. elymoides*: wild, 2.20 mg, field-increased, 2.25 mg, indicating either that seed size for these collections has a strong genetic basis, or that conditions in the agricultural increase field were not sufficiently different from wild-conditions to cause an increase in seed weight. Additional manipulative experiments could investigate the importance of field conditions on seed size. Further, recent evidence suggests that more fine-scaled measures of seed characteristics, such as embryo size, could be more predictive of seed performance than simply size alone (Barak et al. 2018). Even with this uncertainty, we highlighted the larger-seeded sources of both species as the ones that likely have the most beneficial traits.

Finally, of the two commercially available sources tested for each species, there were very few significant differences between them, but for each species, one difference (smaller overall size) suggests that the Vale seed source may perform better than the Toe Jam Creek *E. elymoides* source and that Hanford *P. secunda* (larger seeds) may outperform the Mt. Home cultivar in semi-arid, invaded systems.

## Application and Conclusions

We present this case study as an example of University researchers and Federal practitioners working in direct partnership, combining manager knowledge of remnant native populations with evidence-based strategies to increase restoration success in challenging systems. In the Great Basin, where much of the seed selection and production decisions are centralized outside of particular management areas, this collaboration represents a change from the status quo. As an outcome of this project, wild source populations of the Sonoma collection of *E. elymoides* and the Button Pt. *P. secunda* have been agronomically increased via an IDIQ contract with a private seed grower, resulting in large quantities of locally-collected, agriculturally-increased seeds that have been deployed for projects and experiments within this management area. Projects include two seeding trials implemented in December 2018 and 2019, where seeds from local sources were seeded alongside a more standard commercial mix in three post-fire seedings within the Winnemucca district. These projects are ongoing and will test the performance of these seed sources in realistic management scenarios.

## Acknowledgments

We thank the Great Basin Native Plant Project for funding this work through a research grant to EAL, and thank Shannon Davison, Marenna Disbro, Mariel Boldis, and Dash Hibbard for assistance with root washing and the collection of root measurements and biomass data. The views and opinions of authors expressed herein do not necessarily state or reflect those of the Bureau of Land Management.

## References

Atwater DZ, James JJ, Leger EA (2015) Seedling root traits strongly influence field survival and performance of a common bunchgrass. Basic and Applied Ecology 16:128–140.

Barak RS, Lichtenberger T, Wellman-Houde A, Kramer AT, Larkin DJ (2018) Cracking the case: Seed traits and phylogeny predict time to germination in prairie restoration species. Ecology and Evolution 8:5551–5562.

Barga S, Dilts TE, Leger EA (2017) Climate variability affects the germination strategies exhibited by arid land plants. Oecologia 185:437–452.

Baughman OW, Agneray AC, Forister ML, Kilkenny FF, Espeland EK, Fiegener R, Horning ME, Johnson RC, Kaye TN, Ott J, St. Clair JB, Leger EA (2019) Strong patterns of intraspecific variation and local adaptation in Great Basin plants revealed through a review of 75 years of experiments. Ecology and Evolution 9:6259–6275.

Booth RE, Grime JP (2003) Effects of genetic impoverishment on plant community diversity. Journal of Ecology 91:721–730.

Bower AD, Clair JBS, Erickson V (2014) Generalized provisional seed zones for native plants. Ecological Applications 24:913–919.

Chivers IH, Jones TA, Broadhurst LM, Mott IW, Larson, SR (2016) The merits of artificial selection for the development of restoration-ready plant materials of native perennial grasses. Restoration Ecology 24:174–183.

Davies KW, Boyd CS, Beck JL, Bates JD, Svejcar TJ, Gregg MA (2011) Saving the sagebrush sea: An ecosystem conservation plan for big sagebrush plant communities. Biological Conservation 144:2573–2584.

Enquist CAF, Jackson ST, Garfin ST, Davis FW, Gerber LR, Littell JA, Tank JL, Terando AJ, Wall TU, Halpern B, Hiers JK, Morelli TL, McNie E, Stephenson NL, Williamson MA, Woodhouse CA, Yung L, Brunson MW, Hall KR, Hallett LM, Lawson DM, Moritz MA, Nydick K, Pairis A, Ray AJ, Regan C, Safford HD, Schwartz MW, Shaw MR (2017) Foundations of translational ecology. Frontiers in Ecology and the Environment 15:541–550.

Ferguson SD, Leger EA, Li J, Nowak RS (2015) Natural selection favors root investment in native grasses during restoration of invaded fields. Journal of Arid Environments 116:11–17.

Gibson AL, Espeland EK, Wagner V, Nelson CR (2016) Can local adaptation research in plants inform selection of native plant materials? An analysis of experimental methodologies. Evolutionary Applications 9:1219–1228.

Goergen EM, Leger EA, Espeland EK (2011) Native perennial grasses show evolutionary response to *Bromus tectorum* (cheatgrass) invasion. PLoS ONE 6:e18145.

Haidet M, Olwell P (2015) Seeds of success: A national seed banking program working to achieve long-term conservation goals. Natural Areas Journal 35:165–173.

Hereford J (2009) A quantitative survey of local adaptation and fitness trade-offs. American Naturalist 173:579–588.

Houseal G, Smith DD (2000) Source-identified seed: The Iowa roadside experience. Ecological Restoration 18:173–183.

Johnson RC, Cashman MJ, Vance-Borland K (2012) Genecology and seed zones for Indian ricegrass collected in the southwestern United States. Rangeland Ecology & Management 65:523–532.

Jones TA, Larson SR (2005) Status and use of important native grasses adapted to sagebrush communities. In: Sage-grouse habitat restoration symposium proceedings. RMRS-P-38. US Forest Service, Rocky Mountain Research Station, Fort Collins, CO, USA.

Kiehl K, Kirmer A, Donath TW, Rasran L, Hölzel N (2010) Species introduction in restoration projects – Evaluation of different techniques for the establishment of semi-natural grasslands in Central and Northwestern Europe. Basic and Applied Ecology 11:285–299.

Kulpa SM, Leger EA (2013) Strong natural selection during plant restoration favors an unexpected suite of plant traits. Evolutionary Applications 6:510–523.

Ladouceur E, Jiménez-Alfaro B, Marin M, De Vitis M, Abbandonato H, Iannetta PPM, Bonomi C, Pritchard HW (2018) Native seed supply and the restoration species pool. Conservation Letters 11:e12381.

Leger EA (2013) Annual plants change in size over a century of observations. Global Change Biology 19:2229–2239.

Leger EA (2008) The adaptive value of remnant native plants in invaded communities: An example from the Great Basin. Ecological Applications 18:1226–1235.

Leger EA, Atwater DZ, James JJ (2019) Seed and seedling traits have strong impacts on establishment of a perennial bunchgrass in invaded semi-arid systems. Journal of Applied Ecology 56:1343–1354.

Leger EA, Baughman OW (2015) What seeds to plant in the Great Basin? Comparing traits prioritized in native plant cultivars and releases with those that promote survival in the field. Natural Areas Journal 35:54–68.

Leger EA, Goergen EM (2017) Invasive *Bromus tectorum* alters natural selection in arid systems. Journal of Ecology 105:1509–1520.

Leimu R, Fischer M (2008) A meta-analysis of local adaptation in plants. PLoS ONE 3:e4010.

Vander Mijnsbrugge K, Bischoff A, Smith B (2010) A question of origin: Where and how to collect seed for ecological restoration. Basic and Applied Ecology 11:300–311.

Peppin DL, Fulé PZ, Lynn JC, Mottek-Lucas AL, Hull Sieg C (2010) Market perceptions and opportunities for native plant production on the southern Colorado Plateau. Restoration Ecology 18:113–124.

Pilliod DS, Welty JL, Toevs GR (2017) Seventy-five years of vegetation treatments on public rangelands in the Great Basin of North America. Rangelands 39:1–9.

R Core Team (2019) R: A language and environment for statistical computing.

Riley LE, Steinfeld DE, Winn LA, Lucas SL (2015) Best management practices: An integrated and collaborative approach to native plant restoration on highly disturbed sites. Natural Areas Journal 35:45–53.

Roach DA, Wulff RD (1987) Maternal effects in plants. Annual review of ecology and systematics 18:209–235.

Rowe CLJ, Leger EA (2011) Competitive seedlings and inherited traits: A test of rapid evolution of *Elymus multisetus* (big squirreltail) in response to cheatgrass invasion. Evolutionary Applications 4:485–498.

Stevens AR, Anderson VJ, Fugal R (2014) Competition of squirreltail with cheatgrass at three nitrogen levels. American Journal of Plant Sciences 5:990–996.

