## Supplemental Table 1 for "Using seedling phenotypic traits to select local seed sources for large-scale restoration: methods and outcomes in a Great Basin case study"

**Appendix 1**

In our previous work, earlier emergence has been linked with increased survival in *E. elymoides* (Kulpa & Leger 2013; Leger et al. 2019), though this has not been observed for *P. secunda* (Agneray et al. *unpublished data*). Root length is a factor that has been shown to have very strong association with survival in dry, invaded sites or in drought years, for both *E. elymoides* and *P. secunda* (Atwater et al. 2015; Leger & Goergen 2017; Leger et al. 2019). Increased allocation to roots has been found to be adaptive for *E. elymoides* (e.g. Ferguson et al. 2015; Leger et al. 2019) and *P. secunda* (Leger & Goergen 2017) in some studies, and increased root investment may represent a good strategy for seedlings establishing in dry and invaded areas. Root tips are areas of absorption of nutrients and water and have been associated with increased performance in *Elymus elymoides, E. multisetus* and *Poa secunda* (Atwater et al. 2015, Leger & Goergen 2017). High specific root length, or a high ratio of root length to root mass, usually indicates increased fine root allocation (though it can also indicate differences in root density), and was associated with increased performance of *E. multisetus* (Rowe & Leger 2011) and *E. elymoides* (Leger et al. 2019). In our previous work, we have found multiple lines of evidence that smaller *E. elymoides* individuals were better able to survive under limited-resource conditions (Kulpa & Leger 2013), as well as a general decline in size of multiple Great Basin plants over time (Leger 2013). For *P. secunda*, we did not consider plant size in our consideration of adaptive traits because we lack information about the adaptive nature of relative size in overall small-statured species. Seed size has been a trait with complex effects on field performance in previous experiments. Larger seed size has been shown to increase *E. elymoides* establishment when seed resources are used to grow long roots (Atwater et al. 2015), but may not be beneficial in all field scenarios (Kulpa & Leger 2013). For *P. secunda*, larger seeded sources have been observed to increase survival in some field sites (Agneray et al. *unpublished data*).
